# The cell cycle inhibitor RB is diluted in G1 and contributes to controlling cell size in the mouse liver

**DOI:** 10.1101/2022.06.08.495371

**Authors:** Shuyuan Zhang, Evgeny Zatulovskiy, Julia Arand, Julien Sage, Jan M. Skotheim

## Abstract

Every type of cell in an animal maintains a specific size, which likely contributes to its ability to perform its physiological functions. While some cell size control mechanisms are beginning to be elucidated through studies of cultured cells, it is unclear if and how such mechanisms control cell size in an animal. For example, it was recently shown that RB, the retinoblastoma protein, was diluted by cell growth in G1 to promote size-dependence of the G1/S transition. However, it remains unclear to what extent the RB-dilution mechanism controls cell size in an animal. We therefore examined the contribution of RB-dilution to cell size control in the mouse liver. The RB-dilution model has two requirements. First, manipulations changing RB concentration drive corresponding changes in cell size, and second, the endogenous RB concentration decreases with cell size in G1. We found that both these requirements were met. Genetic perturbations decreasing RB protein concentrations through inducible shRNA expression or through liver-specific *Rb1* knockout reduced hepatocyte size, while perturbations increasing RB protein concentrations in an *Fah^−/−^* mouse model increased hepatocyte size. Moreover, RB concentration decreased in larger G1 hepatocytes while the concentrations of the cell cycle activators Cyclin D1 and E2f1 remained relatively constant. Lastly, we tested how *Rb1* manipulations affected G1/S cell size control in primary hepatocytes using live cell imaging. Loss of *Rb1* weakened cell size control, *i.e.,* reduced the inverse correlation between how much cells grew in G1 and how large they were at birth. Taken together, our results show that an RB-dilution mechanism contributes to cell size control in the mouse liver by linking cell growth to the G1/S transition.

## Introduction

Cell size is important for cell physiology because it determines the size of organelles including the nucleus, spindle, centrosome, mitochondrial network, and vacuole^1–7^. Through this control of organelle size as well as ribosome number, cell size impacts all cellular biosynthetic processes. In addition to this relationship with biosynthesis, cell size is important for the function of diverse cell types. Erythrocytes are tiny because they must migrate through tight spaces, while macrophages must be large enough to engulf pathogens. The importance of cell size for diverse cell types is further reflected in the fact that the size of cells of a given type is remarkably uniform and deviations from the typical cell size are often associated with disease states. For example, the coefficient of variation of the size distribution of red blood cells is used as a common diagnostic parameter in comprehensive blood tests, and many cancers, such as small-cell lung cancer, are characterized by altered cell size^8–14^.

To control their size to be within a target range, proliferating cells can either regulate their growth such that larger cells grow slower, or they can link cell growth to progression through the cell division cycle^15–19^. Studies in cultured cells have now begun to elucidate some of the molecular mechanisms that are responsible. One model proposes that cell growth triggers the G1/S transition by diluting the cell cycle inhibitor RB, the retinoblastoma protein^20^. The amount of dilution required, and the target cell size, can then be modulated by the activity of the cyclin D-Cdk4 kinase complex^21^. A second model proposes that small cell size activates the p38 stress kinase, which, in turn, inhibits cell division^22^. Taken together, this recent work in the field is starting to uncover a series of molecular mechanisms through which cell growth triggers the G1/S transition to control cell size. However, these cell size control mechanisms are all based on *in vitro* cell culture studies, and their *in vivo* relevance remains unclear.

Here, we aimed to test the *in vivo* relevance of one cell size control model, namely the RB-dilution model, in the liver of genetically engineered mice. Consistent with the RB-dilution model, increasing or decreasing RB protein concentrations in hepatocytes *in vivo* caused an increase or decrease in their cell size, respectively. In addition, we found that RB protein concentration decreased with cell size in G1 phase, consistent with RB concentration being a cell size sensor. Finally, we found that RB deletion reduced the ability of primary hepatocytes to control their size when grown in culture. More specifically, we found that deletion of *Rb1* reduced the inverse correlation between how large hepatocytes were at birth and how much they grew in G1. Taken together, our results suggest that the RB-dilution mechanism plays an important role in controlling hepatocyte cell size.

## Results

### Decreasing RB protein concentration in mouse liver reduces hepatocyte size

If the RB-dilution model applies to the mouse liver, reducing RB protein concentration would reduce hepatocyte size. To test this, we utilized two different mouse models to reduce RB protein concentration. The first model harbors a Doxycycline-inducible transgene broadly expressing an shRNA against *Rb1 (Rosa-rtTA; TRE-shRb1* mouse^23^). We gave Doxycycline water (1g/L) from birth to 4 weeks old to *Rosa-rtTA* (control) or *Rosa-rtTA; TRE-shRb1* mice from the same litter (Figure 1A). Then, the hepatocytes were isolated from these mice to check for knockdown efficiency (Figure 1B) and to measure their cell size distributions (Figure 1C). Since the liver is a highly polyploid organ^24^, and polyploid cells are bigger than diploid cells, we sorted the isolated hepatocytes based on viability and on ploidy before size measurements were made in order to eliminate the confounding effect of ploidy on size (Figure 1C). We found that within the same ploidy, the hepatocyte size in *Rb1* knockdown mice was smaller than that in the control mice, supporting the RB-dilution model.

**Figure 1.**
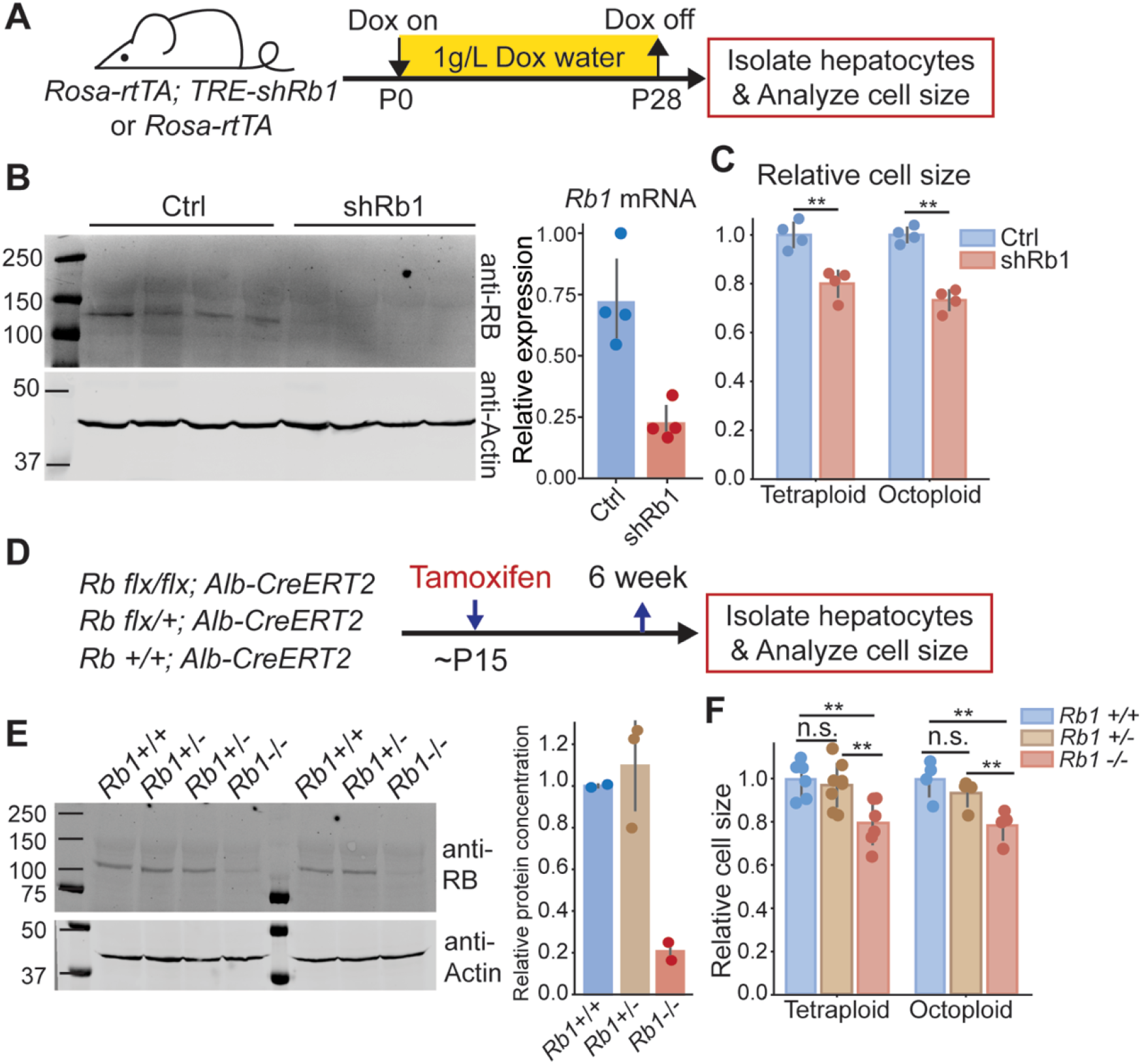
Decreasing Rb1 protein concentration in mouse liver reduces hepatocyte size. A. Schematic of experimental timeline performed using *Rosa-rtTA; TRE-shRb1* mice. B. Immunoblot and qPCR analysis following shRb1 induction in mouse liver (n = 4). Mice were harvested after 4 weeks of Dox treatment (1g/L) that started at birth. C. Cell size measurement of the hepatocytes isolated from *Rosa-rtTA* (Ctrl) and *Rosa-rtTA; TRE-shRb1* (shRb1) mice by Coulter counter. D. Schematic of experimental timeline performed using *Rb1^flx/flx^; Alb-CreERT2* mice. E. Immunoblot shows Rb1 protein concentrations in the livers of *Rb1^+/+^, Rb1^+/-^*, and *Rb1^−/−^* mice. Mice were harvested 6 weeks after Tamoxifen treatment. The right panel shows the quantification of protein concentration. RB protein concentration was normalized using actin. F. Cell size measurements were made using a Coulter counter to examine hepatocytes isolated from *Rb1^+/+^, Rb1^+/-^*, and *Rb1^−/−^* mice at 6 weeks after Tamoxifen treatment. Each dot represents data from one mouse. The error bars indicate standard deviation. ** *P*<0.01.

To further test the RB-dilution model in the liver, we used a second mouse model that also reduced RB protein concentrations, but only in the liver. This mouse model contains an *Rb1^flx/flx^* allele and expresses the Cre recombinase only in the liver through an *Alb-CreERT2* allele. By injecting Tamoxifen into these mice, the *Rb1* gene can be knocked out in a liver-specific manner. We generated *Rb1^−/−^, Rb1^-/+^, and Rb1^+/+^* mice from the same litter by administering Tamoxifen to these mice when they were 15 days old (Figure 1D). At 6 weeks of age, we harvested the livers and examined the hepatocyte size of these mice. We note that the *Rb1^+/-^* mice did not have significantly reduced RB protein concentrations compared to the *Rb1^+/+^* mice (Figure 1E). Consistent with this similar RB protein expression, we found that hepatocyte size in *Rb1^+/-^* mice was not significantly different from that in *Rb1^+/+^* mice (Figure 1F). In contrast, hepatocyte size was significantly smaller in *Rb1^−/−^* mice compared to *Rb1^+/+^* and *Rb1^+/-^* mice, which again supports a role for RB in controlling cell size in the mouse liver.

### Increasing RB concentration in mouse liver increases hepatocyte cell size

After finding that decreasing RB concentration decreases cell size, we sought to test if increasing RB protein concentration increases hepatocyte cell size. To overexpress RB, we utilized the *Fah^−/−^* mouse system^25^. Deletion of the *Fah* gene causes toxin accumulation in hepatocytes that will eventually lead to hepatocyte death. Toxin accumulation is prevented in *Fah^−/−^* mice by treating them with NTBC (2-(2-nitro-4-trifluoromethylbenzoyl)-1,3- cyclohexanedione)^26^. When NTBC is withdrawn, cells expressing exogenous *Fah*, introduced by injecting *Fah*+ transposons, will clonally expand to repopulate the injured liver (Figure 2A)^27^. Importantly, other genetic elements, such as transgenes, can be added to the *Fah* transposon so that they are co-integrated into some hepatocyte genomes. This system provides a versatile platform for genetic manipulations in the liver. To increase RB expression, we delivered an *Fah-P2A-Rb1* transposon and the SB100 transposase into *Fah^−/−^* mice via hydrodynamic transfection through the tail vein. As a control, we injected the transposon with *Fah-P2A-mCherry* (Figure 2B). 6 weeks after injection, the liver was almost fully repopulated by the cells containing *Fah* transposons, and there was an appreciable level of Rb1 protein overexpression (Figure 2C, D). Consistent with the RB-dilution model, overexpression of RB resulted in bigger hepatocytes (Figure 2E).

**Figure 2.**
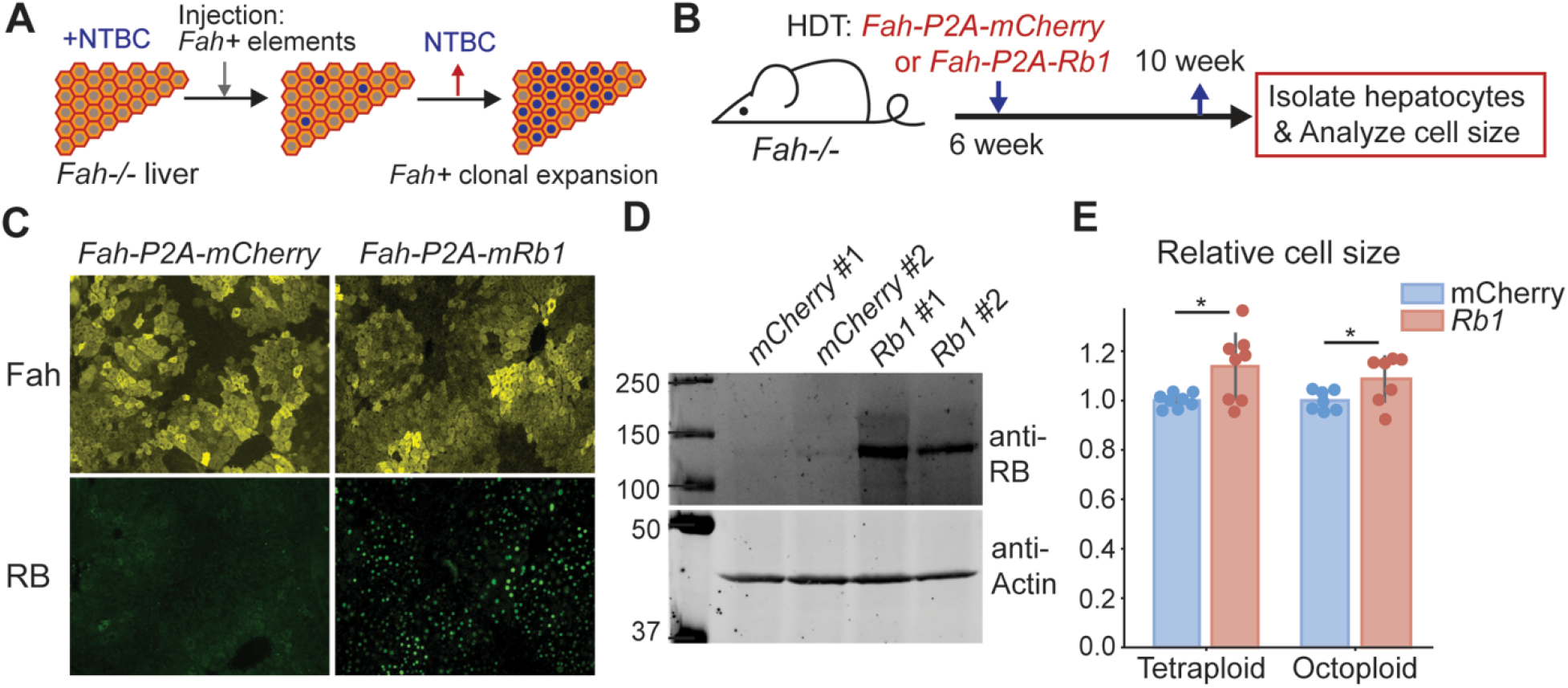
Increasing Rb1 concentration in mouse liver increases hepatocyte cell size. A. Schematic of *Fah-/-* system as a platform for genetic manipulation of hepatocytes in the liver. B. Schematic of experimental timeline. *Fah^−/−^* mice were injected with *Fah* transposons as indicated, and their livers were harvested 4 weeks after the injection. C-D. Immunofluorescence staining (C) and immunoblots (D) were performed to measure Fah and RB expression. E. Cell size measurement of the hepatocytes isolated from *Fah^−/−^* mice injected with *Fah* transposons were made with a Coulter counter. Each dot represents data from one mouse. The error bars indicate standard deviation. * *P*<0.05.

In addition to RB dosage affecting cell size in cultured cells, it was also reported that the activity of the upstream kinase cyclin D-Cdk4/6 affected cell size^21^. Lower Cdk4/6 activity resulted in larger cells, while higher activity reduced cell size. This can be integrated with the RB-dilution model since cyclin D-Cdk4/6-dependent RB phosphorylation contributes to RB inactivation. This means that the higher the Cdk4/6 activity, the more RB is phosphorylated and inactivated, and the less it needs to be diluted^28^. To test for the effect of Cdk4/6 activity on hepatocyte cell size, we used Palbociclib to inhibit Cdk4/6 activity during liver regeneration following a partial hepatectomy (Supplementary Figure 1A). In this experiment, the partial hepatectomy induces massive hepatocyte growth and proliferation during a short time window that can be monitored experimentally. More specifically, following partial hepatectomy, mice were treated with Palbociclib (80mg/kg) for 5 consecutive days (Supplementary Figure 1A). We isolated hepatocytes from these mice at different time points and measured cell size. Consistent with the observations in cultured cells that the Cdk4/6-RB pathway controls cell size, Palbociclib treatment increased hepatocyte size following a partial hepatectomy (Supplementary Figure 1B).

### RB protein is diluted in G1 in primary hepatocytes

So far, our data indicate that manipulations of RB protein concentrations *in vivo* changed hepatocyte size as the RB-dilution model predicted. However, these deletion and overexpression experiments cannot determine if RB protein concentration functions as a cell size sensor. For RB to be a cell size sensor responsible for cell size control, not only must its concentration affect cell size, but its concentration must also change with cell size. In other words, RB must be diluted in G1 for the model to work.

To test if RB is diluted in G1, we sought to measure the amounts of RB protein and cell size in single cells. However, detecting RB protein in single proliferating cells from the mouse liver tissue is challenging for several reasons. First, immunofluorescence measurements are challenging due to RB’s low expression level and the high autofluorescence background of hepatocytes. Second, RB-dilution would control cell size by coordinating cell growth with the G1/S transition only in proliferating cells, who represent only a minority of adult hepatocytes^29,30^. To circumvent these challenges, we sought to specifically measure RB protein concentrations in proliferating isolated primary hepatocytes. To do this, we used a 2D primary hepatocyte culture protocol^31,32^ (Figure 3A). In these cultures, primary hepatocytes actively enter the cell cycle, and we can detect RB protein using immunfluorescence because the autofluorescence is significantly reduced in the isolated hepatocytes compared to those in the mouse liver (Figure 3C).

**Figure 3.**
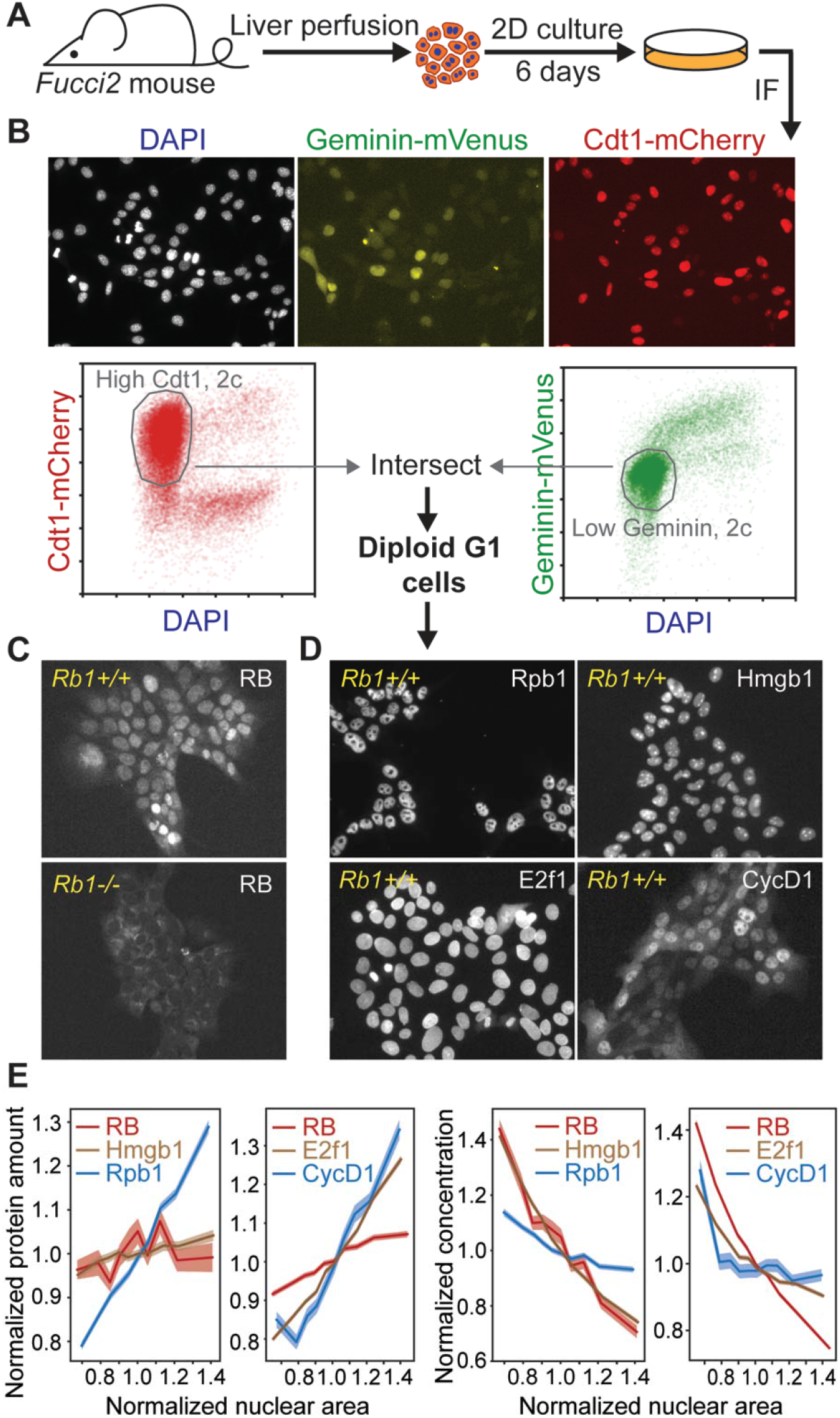
RB protein is diluted in G1 in primary hepatocytes. A. Schematic of experiment examining Rb1 protein concentrations in single primary hepatocytes in 2D cultures. B. Representative immunofluorescence images of primary hepatocytes isolated from *Fucci2* mice and illustration of gating procedure used to isolate G1 cells. C. Immunofluorescence staining of RB in *Fucci2* primary hepatocytes. The *Rb1^−/−^* hepatocytes were used as a negative control for RB antibody staining. D. Immunofluorescence staining of Rpb1, Hmgb1, CycD1, and E2f1 in *Fucci2* primary hepatocytes grown in 2D culture. E. Nuclear area of G1 hepatocytes plotted against the indicated protein amount (left panels) or concentration (right panels). The shaded area indicates 95% confidence interval.

To measure the RB protein concentrations in hepatocytes, we performed an immunofluorescence staining on 2D hepatocyte cultures isolated from *Fucci2* mice expressing G1 and S/G2 cell cycle phase reporters^33^ (Figure 3A). Based on their DNA content, Cdt1-mCherry signal, and Geminin-Venus signal, we determined which cells in the population were in G1 phase (Figure 3B). We then examined the relationship between target protein amount and cell size, which was estimated using nuclear area as a proxy for cell size^34^. We also measured the RNA Polymerase II subunit Rpb1, whose concentration was expected to stay constant, and Hmgb1, which has been found to be diluted by cell growth similarly to RB in other cell lines (Figure 3D)^35^. We found that unlike Rpb1, the total amount of RB protein did not increase in proportion to cell size, but behaved more similarly to the anticipated subscaling protein Hmgb1 (Figure 3E). We also examined amounts of two key activators of G1/S transition, Cyclin D1 and E2f1 (Figure 3D). While RB protein amounts remained relatively unchanged as cells became larger in G1, Cyclin D1 and E2f1 protein amounts increased with cell size (Figure 3E), suggesting that the activator (Cyclin D1, E2f1) to inhibitor (RB) ratio is increasing as cells grow in G1. Our measurements of RB concentration as a function of nuclear area demonstrate that RB is diluted as cells grow in G1. Taken together, our immunofluorescence measurements of RB concentration in proliferating hepatocytes are consistent with RB operating as a cell size sensor in G1.

### Knocking out *Rb1* weakens G1/S cell size control in primary hepatocytes

At its most fundamental level, cell size control requires cells that are born smaller grow more during the cell division cycle than cells that are born larger. Size control thus results in more similarly sized cells following a division cycle. The degree to which cells control their size can therefore be quantified by the inverse correlation between cell size at birth and the amount of cell growth. Therefore, the strength of cell size control can be evaluated by the slope of the linear fit between the cell size at birth and the amount of cell growth over the cell cycle^9,36^. Perfect size control would yield a slope value of −1, and if the amount of growth was independent of the birth size, the slope would be 0. If RB contributed to cell size control, its deletion should reduce the degree of inverse correlation between cell size at birth and the amount of cell growth in the subsequent cell cycle^20^.

To test the extent to which RB was responsible for cell size control in G1, we sought to measure cell size and cell cycle phase durations in proliferating hepatocytes that we tracked through cell divisions. To do this, we performed live cell imaging on the 2D cultured hepatocytes isolated from *Rb1^flx/flx^; Rosa26-CreERT2; Fucci2* mice. 4 days before imaging, cells were treated with 4- Hydroxytamoxifen (4-OHT) to delete the *Rb1* gene (Figure 4A). We used the inflection point of the Geminin-Venus S/G2 reporter signal and the maximum point of the Cdt1-mCherry G1 reporter signal to determine the G1/S transition time at the midpoint between these two events (Figure 4B). We again used the nuclear area as a proxy for cell size. Consistent with RB contributing to cell size control, we found that the *Rb1^−/−^* hepatocytes exhibited smaller size and shorter G1 length (Figure 4C). Moreover, the correlation between birth nuclear area and the amount of growth in G1 was reduced in *Rb1^−/−^* hepatocytes compared to *Rb1^+/+^* hepatocytes (Figure 4D, E, Supplementary Figure 1C, D). We note that in each experiment, we used hepatocytes from different mice, but that experimental and control hepatocytes for each replicate were derived from the same mouse. Taken together, our data show that knocking out *Rb1* weakened G1/S cell size control in hepatocytes, consistent with the RB-dilution model.

**Figure 4.**
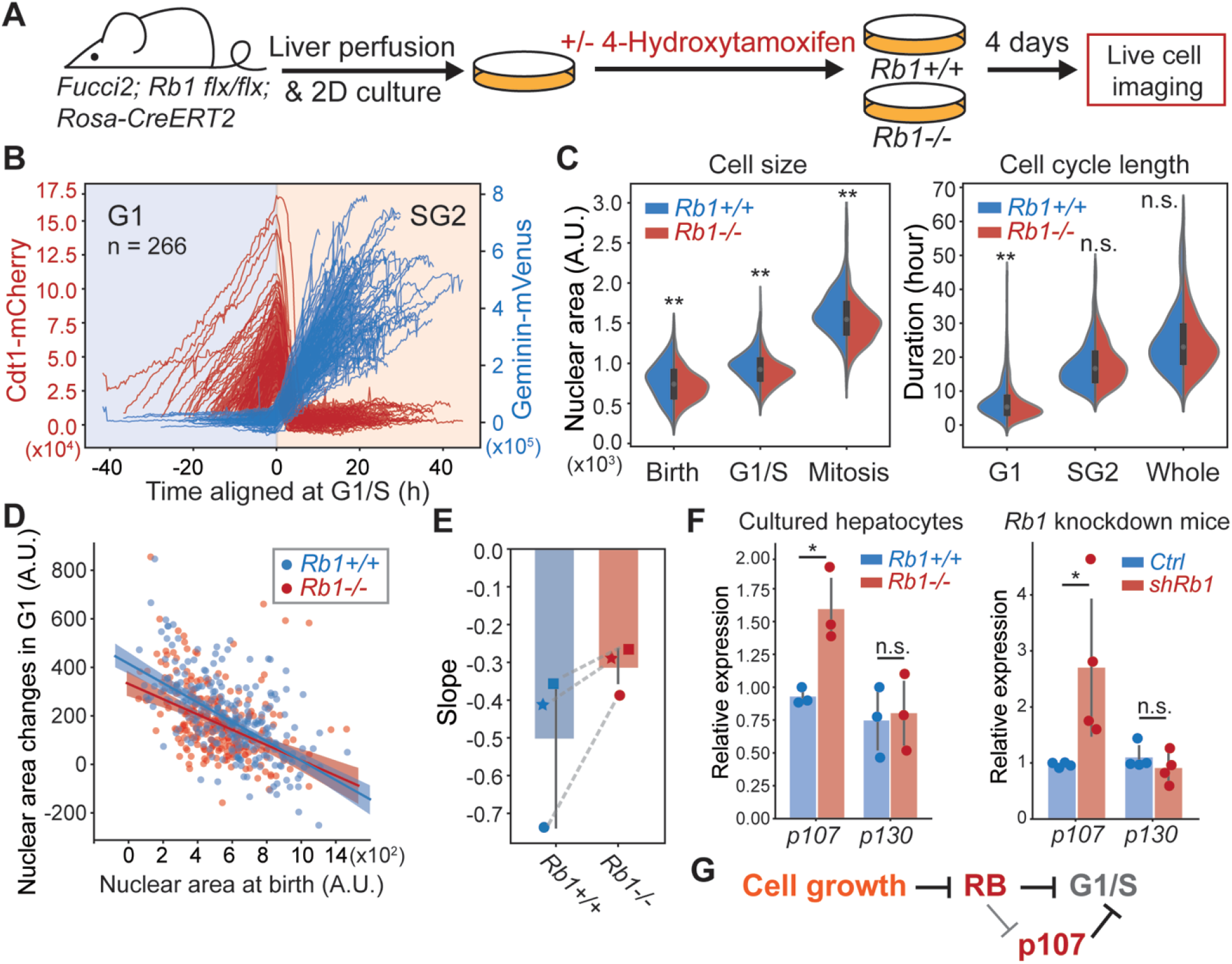
Deletion of *Rb1* weakens G1/S cell size control in primary hepatocytes. A. Schematic of the experiment used to examine the role of *Rb1* in hepatocyte cell size control. B. Live cell imaging traces of the Cdt1-mCherry and Geminin-Venus cell cycle reporters in *Fucci2* hepatocytes grown in 2D culture. The G1/S transition is determined using the inflection point of the Geminin-Venus signal and the point of maximum Cdt1-mCherry signal. C. Cell size and cell cycle statistics of *Rb1^+/+^* and *Rb1^−/−^* hepatocytes. Cell size is approximated using nuclear area. D. The correlations between nuclear area at birth and the nuclear area changes during G1 in *Rb1^+/+^* and *Rb1^−/−^* hepatocytes. The shaded area indicates the 95% confidence interval. E. The slope values of the linear relationship in (D) are plotted from three independent biological replicates. Each experiment used hepatocytes from different mice, and *Rb1^+/+^* and *Rb1^−/−^* hepatocytes for each replicate experiment were derived from the same mouse. Dashed lines link experiments performed using hepatocytes derived from the same mouse. *F. p107* mRNA measured by RT-qPCR from *Rb1^+/+^* and *Rb1^−/−^* 2D cultured primary hepatocytes (left panel) and *Rosa-rtTA; TRE-shRb1* mice (right panel). The error bars indicate standard deviation. * *P*<0.05, ** *P*<0.01. G. Model indicating multiple mechanisms through which RB regulates cell size at the G1/S transition.

While *Rb1* deletion weakened G1/S size control, it did not remove it completely as the slope between nuclear area at birth and G1 growth amount was not 0 (Figure 4D). It was previously shown that *Rb1* deletion resulted in the upregulation of the related protein p107, which can compensate for *Rb1* loss^37^. Therefore, the partial loss of size control in *Rb1^−/−^* cells might be explained by p107 upregulation because p107 protein was previously found to be similarly diluted as RB^20^. Moreover, the combined deletion of *Rb1* and *p107* genes in mouse embryonic fibroblasts resulted in much smaller cells than either of the single deletions^37,38^. To test if *p107* was upregulated in *Rb1^−/−^* hepatocytes, we measured the mRNA levels of *p107* in *Rb1^−/−^* 2D cultured hepatocytes, as well as in the *Rosa-rtTA; TRE-shRb1* mouse livers. Indeed, *p107* mRNA was increased in *Rb1^−/−^* and *Rb1* knock down cells (Figure 4F), supporting our hypothesis that *p107* is upregulated to compensate for *Rb1* loss and contributes to G1/S size control when *Rb1* was deleted.

## Discussion

Cell size is fundamental to cell physiology, yet we are only now beginning to understand the molecular mechanisms that control size through linking cell growth to division. Cell growth was found to trigger cell division by diluting a cell cycle inhibitor in budding yeast and *Arabidopsis* plant cells^35,39–42^. This general principle of inhibitor-dilution was also found in human cells grown in culture, who diluted the cell cycle inhibitor RB in G1 to trigger the G1/S transition^20^. This RB-dilution mechanism can be modulated by cyclin D-Cdk4/6 activity, which inactivates RB^21^. However, it was not clear whether the RB-dilution mechanism was physiologically relevant to cells growing in tissues *in vivo.* This is because of the large differences between the conditions cells experience when growing in a 2D *in vitro* culture compared to a 3D tissue environment. In tissues, cells send and receive many more paracrine and endocrine signals, have multiple cellcell contacts, experience dynamic forces and pressures, and experience an environment with very different nutrients and growth factors. In addition to these differences in culture and tissue environments, size control models developed in culture are cell autonomous and do not take into account the possibility that tissue level processes could be used to sense and control the size of neighboring cells.

Due to the significant differences of *in vitro* and *in vivo* growth conditions, it is crucial to test molecular models developed using cell culture in animal contexts. We therefore sought to test the RB-dilution model *in vivo* in the mouse liver. The RB-dilution model has two main requirements. First, that changing the concentration of RB should have an effect on the average cell size, and second, that the concentration of endogenous RB should reflect cell size. Both of these requirements were satisfied in mouse hepatocytes. By using genetically engineered mouse models to manipulate RB concentrations, we found hepatocyte size was correlated with RB protein concentrations in mouse liver. Moreover, the endogenous RB protein was diluted in larger G1 cells. Finally, we examined cell size control in proliferating isolated primary hepatocytes transferred into culture. Loss of *Rb1* weakened G1/S cell size control, *i.e.,* reduced the correlation between how much cells grew in G1 and how large they were at birth. Taken together, these results support a role for the RB-dilution model in regulating hepatocyte cell size. As previous work showed similar mechanisms operate in yeast and plants, it is now clear that examples of the inhibitor-dilution class of mechanisms can be found across the tree of life.

In addition to being broadly applicable, inhibitor dilution models can easily be modified to link cell size to cell cycle progression in polyploid cells, which are common in mouse liver. Indeed, polyploidy is a common feature of mammalian livers, where ploidy is generally proportional to cell size^43^. The RB-dilution model is compatible with hepatocyte polyploidization, which is predominantly mediated by cytokinesis failure^44,45^. This is because the link between cell growth and cell cycle progression through the dilution of a cell cycle inhibitor does not actually require division. RB may be diluted in G1 so that S phase is triggered. The, RB accumulates through S/G2 and cells can return to a subsequent stable tetraploid G1 phase without division. Crucially, the RB-dilution model is based on protein concentrations, which will be the same in two diploid G1 cells (if division was successful) as in one tetraploid G1 cell (if division was unsuccessful). Inhibitor dilution thus potentially provides an elegant mechanism to explain the repeated growth requirements for DNA synthesis and the large size of polyploid eukaryotic cells.

While our data here support a role for RB-dilution in controlling cell size in the mouse liver, they also indicate that this is unlikely to be the only size control mechanism working in this context. The presence of compensatory mechanisms can be inferred from the fact that *Rb1* deletion and over-expression produce modest changes in cell size (Figures 1–2). Moreover, *Rb1* deletion in primary hepatocytes grown in culture only partially eliminated the correlation between growth and progression through G1 (Figure 4). One possible explanation for these results is that RB regulates the expression of the related cell cycle inhibitor p107^37,38,46,47^. Consistent with prior work, we found that deletion of *Rb1* In hepatocytes strongly increased p107 expression. This regulation of *p107* by RB could buffer cell size against changes in RB protein concentrations. Further investigation into this mechanism is needed, which can be achieved by incorporating *p107* knockout in the *Rb1* knockout background.

In addition to p107 regulation buffering the effect of *Rb1* deletion, there could be additional size-control mechanisms. In yeast, Whi5-dilution is one of several mechanisms that operate based on size-dependent gene expression^48,49^. In this model, a series of cell cycle activators generally increase in concentration as cells grow, while cell cycle inhibitors decrease in concentration. The balance of activators and inhibitors then sets cell size. Removal of any one component would serve to shift the mean cell size, but not remove the ability of the cell to control size around the new setpoint. Supporting the applicability of this multiple size-scaling mechanism model to mammalian cells, recent proteomics analyses found that cell size-dependent concentration changes were widespread across the proteome of human cells grown in culture^35,50^. Moreover, beyond such size-scaling concentration-based mechanisms, there may be additional mechanisms operating by other mechanistic principles yet to be discovered. Much more work remains to be done to identify the mechanistic links between cell growth and division.

As a first step, we anticipate such work will need to be done in yeast and cultured cells. While this work will need to be validated *in vivo,* our results here, indicating that the RB-dilution model contributes to controlling cell size in hepatocytes, should serve as encouragement for future studies *in vitro* and their application *in vivo.*

## Acknowledgements

This work was supported by the National Institutes of Health (R01 DK128578, P01 CA254867). We thank Roel Nusse and Jinghua Li for assistance in establishing the protocol to culture primary hepatocytes in 2D, Thuyen Nguyen for maintaining the mouse colonies in the Sage lab, Dr. Marcus Grompe for sharing the *Fah^−/−^* mice, and the members of the Skotheim laboratory for discussions and feedback on the work.

## Materials and Methods

### Mice

All mice were handled in accordance with the guidelines of the Institutional Animal Care and Use Committee at Stanford University. The *Rosa-rtTA; TRE-shRb1* mice were described before^23^. *Rosa-rtTA* or *Rosa-rtTA; TRE-shRb1* mice were given Doxycyclin water (RPI, 1g/L) from birth to when they were 4 weeks old. At 6 weeks old, the mice were sacrificed to isolate primary hepatocytes. *Alb-CreERT2* mice were a kind gift of Dr. Pierre Chambon^51^, and *Rb1^flx/flx^; p107^−/−^; p130^flx/flx^*; *Alb-CreERT2* were described before^52^. We bred *Rb1^flx/flx^; p107^−/−^; p130^flx/flx^; Alb-CreERT2* mice to *Rb1^flx/flx^* mice to generate *Rb1^flx/flx^; Alb-CreERT2, Rb1^flx/+^; Alb-CreERT2, and Rb1^+/+^: Alb-CreERT2* mice from the same cohort. Then, tamoxifen (Sigma, 50mg/kg, 5 consecutive doses) was given to these mice when they were ~15 days old through i.p. injection. At 6 weeks old, the mice were sacrificed to isolate primary hepatocytes. The *Fah^−/−^* mice were kindly shared by Dr. Markus Grompe’s group at Oregon Health & Science University. At 6 weeks old, the *Fah^−/−^* mice were subjected to hydrodynamical transfection through their tail vein. The DNA plasmids we injected contain a transposon and transposase. The plasmid backbone was kindly provided by Dr. Hao Zhu’s lab at UT Southwestern Medical Center. We modified the transposons to express the *Fah* gene and mCherry or Rb1. The transposase was SB100. 6 weeks after the injection, the mice were sacrificed to isolate primary hepatocytes. The *Fucci2* mice^33^ were kindly shared by Dr. Valentina Greco’s lab at Yale University. The *Fucci2* mice contained the transgene expressing Cdt1-mCherry and Geminin-Venus from the *ROSA26* locus.

### Partial hepatectomy and Palbociclib treatment

2/3^rd^ partial hepatectomy was performed on 8-week-old male CD1 mice (JAX). The procedure followed an established protocol^53^. At day 0 to day 5 after the surgery, mice were injected with either vehicle or Palbociclib HCl (80mg/kg). Palbociclib HCl was dissolved in 50mmol/L sodium lactate (pH 4) at a concentration of 10μg/mL under a fume hood. Then, the drug was given to mice at a dosage of 80mg/kg via oral gavage for 5 consecutive days. On day 0, 1, 2, 3, 4, 5 after surgery, one mouse from each treatment was sacrificed to isolate primary hepatocytes.

### Primary hepatocyte isolation and size measurement

Primary hepatocytes were isolated by two-step collagenase perfusion^54^. Liver perfusion medium (Thermo Fisher Scientific, 17701038), liver digest medium (Thermo Fisher Scientific, 17703034) and hepatocyte wash medium (Thermo Fisher Scientific, 17704024) were used. After isolation, cells were washed with PBS once and stained with Zombie Green (Biolegend, 423111) at room temperature for 15 minutes to mark dead cells. After washing with cold PBS twice, cells were fixed in 4% PFA (Thermo Fisher Scientific, 28906) at 37°C for 15 minutes. Then, cells were washed with PBS 3 times, subjected to DNA staining using Hoechst (Thermo Fisher Scientific, 62249), and sorted on a Flow Cytometer (BD FACSAria II) based on their live/dead staining and their DNA staining. Only live (without Zombie Green signal) hepatocytes with same ploidy were sorted. 10000-20000 cells were sorted for each population. After sorting, the cell size distributions for each population were measured using a Z2 Coulter counter (Beckman).

### Primary hepatocyte 2D culture

The protocol for 2D culture of primary hepatocytes was kindly shared by Yinhua Jin from Dr. Roeland Nusse’s lab at Stanford University^31,32^. Briefly, primary hepatocytes from *Rb1^flx/flx^; Rosa26-CreER2; Fucci2* mouse were isolated by two-step collagenase perfusion. After isolation, cells were washed 3 times with hepatocyte wash medium (Thermo Fisher Scientific, 17704024). Cells were then plated in a 6-well plate precoated with collagen I (50μg/mL) at a density of 200,000 per well. The culture medium contained 3μM CHIR99021 (Peprotech), 25ng/mL EGF (Peprotech), 50ng/mL HGF (Peprotech), and 100ng/mL TNFa (Peprotech) in Basal medium. The Basal medium contained William’s E medium (GIBCO), 1% Glutamax (GIBCO), 1% Non-Essential Amino Acids (GIBCO), 1% Penicillin/streptomycin (GIBCO), 0.2% normocin (Invitrogen), 2% B27 (GIBCO), 1% N2 supplement (GIBCO), 2% FBS (Corning), 10mM nicotinamide (Sigma), 1.25mM N-acetylcysteine (Sigma), 10μM Y27632 (Peprotech) and 1μM A83-01 (Tocris). The culture medium was refreshed every other day. Cells were passaged via trypsinization using TrypLE (Thermo-Fisher).

### Live cell imaging and analysis

The cultured Fucci2 hepatocytes were seeded on collagen coated 35-mm glass-bottom dishes (MatTek) one day before imaging. Then, the cells were transferred to a Zeiss Axio Observer Z1 microscope equipped with an incubation chamber and imaged for 48 hours at 37°C and 5% CO_2_. Brightfield and fluorescence images were collected from *Rb1^+/+/^* and *Rb1^−/−^ Fucci2* hepatocytes at multiple positions every 20 minutes using an automated stage controlled by the Micro-Manager software. We used a Zyla 5.5 sCMOS camera and an A-plan 10x/0.25NA Ph1 objective. For each cell, the transition from G1 to S phase was taken as the midpoint between when the inflection point of the Geminin-Venus signal and the point of maximum Cdt1-mCherry signal occurred. Cell size was approximated using the area of the nucleus^34^.

### Immunoblots

Mouse liver tissues or isolated primary hepatocyte pellets were homogenized and lysed in RIPA buffer (Thermo Scientific) supplemented with protease inhibitors and phosphatase inhibitors. Lysates were separated on 8% SDS-PAGE gels and transferred to nitrocellulose membranes. Membranes were then blocked with SuperBlock™ (TBS) Blocking Buffer (Thermo Fisher Scientific) and incubated overnight at 4°C with primary antibodies in 3% BSA solution in PBS. The primary antibodies were detected using the fluorescently labeled secondary antibodies IRDye® 680LT Goat anti-Mouse IgG (LI-COR 926-68020) and IRDye® 800CW Goat anti-Rabbit IgG (LI-COR 926-32211). Membranes were imaged on a LI-COR Odyssey CLx and analyzed with LI-COR Image Studio software. The following antibodies were used: anti-β-Actin (Sigma, A2103, 1:2000), Anti-Rb (Santa cruz, sc-74570, 1:500).

### Immunofluorescence staining

*Fucci2* hepatocytes were seeded on a 35-mm collagen-coated glass-bottom dish (MatTek) one day before immunofluorescence staining and analysis. For the staining, cells were fixed with 4% paraformaldehyde for 20 minutes at room temperature, permeabilized with 0.5% Triton™ X-100 (Sigma-Aldrich) for 15 minutes at room temperature, and then blocked with 3% BSA in PBS.

Then, the cells were incubated with primary antibodies overnight at 4°C. After 3 washes with PBS, the cells were incubated with Alexa Fluor 647 conjugated goat anti-mouse secondary antibodies (Invitrogen, A32728) at 1:1000 for 1 hour at room temperature. After 3 washes with PBS, cells were incubated with 500 nM DAPI for 30 minutes at room temperature before imaging. The primary antibodies used for immunofluorescence were Anti-Rb (Santa cruz, sc- 74570, 1:100), Anti-Rpb1-CTD (Abcam, ab252854, 1:250), Anti-HMGB1 (Abcam, ab79823, 1:250), Anti-E2F1 (Santa cruz, sc-251, 1:100), and Anti-Ccnd1 (Abcam, ab16663, 1:250). The cells were imaged using a Zeiss Axio Observer Z1 microscope with an A-plan 10x/0.25NA objective.

### Image analysis

For microscopy data quantification, cell nuclei were segmented using either the mCherry channel for cells in G1 that express Cdt1-mCherry or the GFP channel for cells in S/G2 that express Geminin-Venus. Segmentation was performed using the Fiji plugin StarDist, which is a deep-learning tool for segmenting nuclei in images that are difficult to segment using thresholding-based methods. The total pixel intensities within the segmented masks in each channel were recorded, and each object’s background was subtracted based on the median intensity of the image. G1 cells were selected based on their Cdt1-mCherry and Geminin-Venus signals. The cells that had high Cdt1-mCherry and low Geminin-Venus were selected as G1 cells. When analyzing the scaling behavior of immunostained proteins, nuclear area was used as a proxy for cell size^7^.

### RNA extraction and qRT-PCR

Total RNA was isolated using Direct-zol RNA Miniprep kit (Zymo Research). For qRT-PCR, cDNA synthesis was performed with 1μg of total RNA using an iScript Reverse Transcription Kit (Biorad). qPCR reactions were made with the 2x SYBR Green Master Mix (Biorad). Gene expression levels were measured using the ΔΔCt method.

### Statistical analysis

The data in most figure panels reflect multiple experiments performed on different days using mice derived from different litters. For comparison between groups, we conducted the hypothesis test using the two-tailed student’s t-test. Statistical significance is displayed as p < 0.05 (*) or p < 0.01 (**) unless specified otherwise.

## Supplementary Material

**Figure S1:**
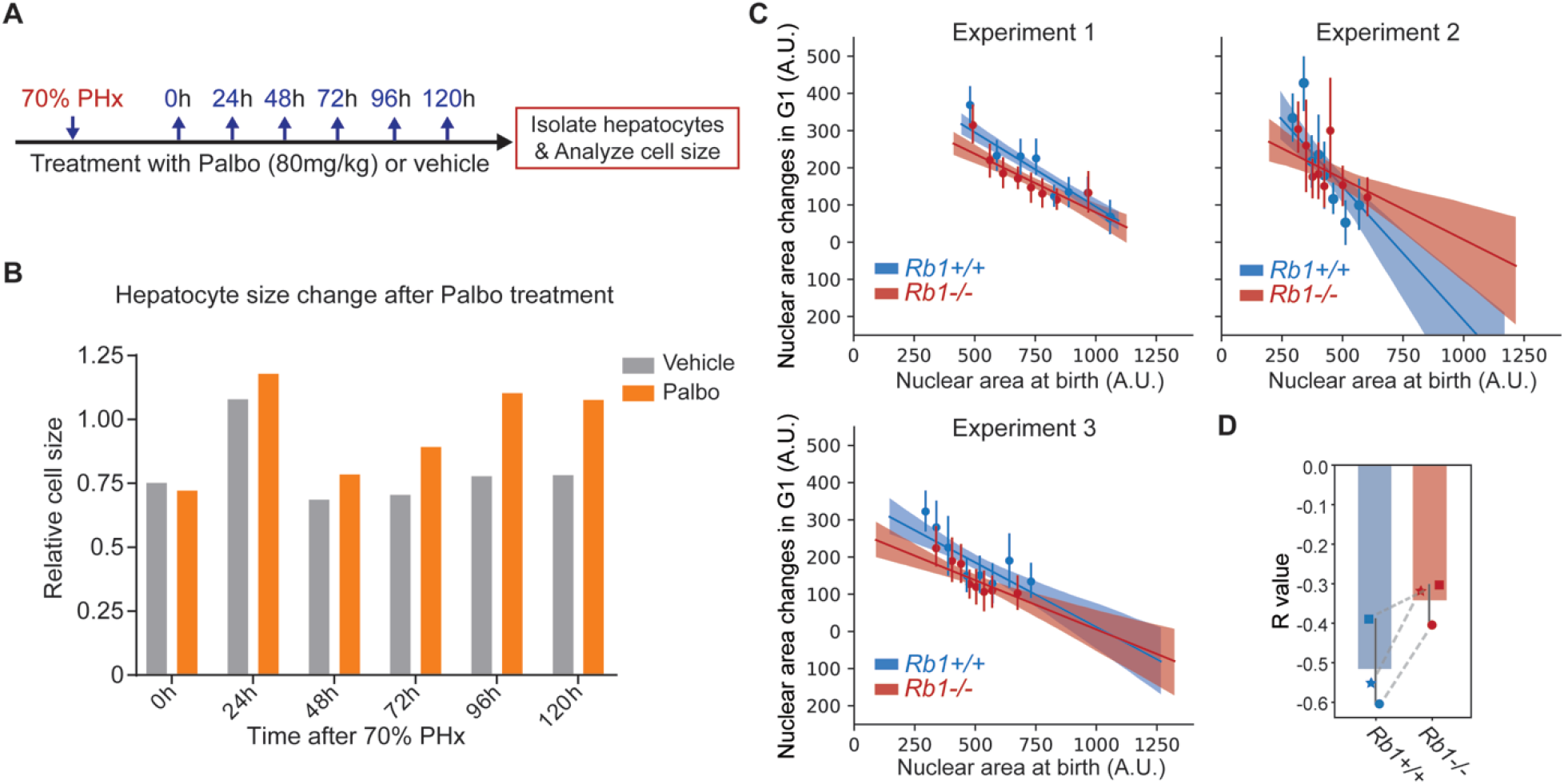
Cell size analysis following hepatectomy, and single cell time lapse imaging data for each individual experiment. A. Schematic of the partial hepatectomy (PHx) experiment examining the effect of Palbociclib treatment. B. Coulter counter cell size measurements of hepatocytes isolated from mice at the indicated time points following surgery. C. The correlations between nuclear area at birth and the nuclear area changes during G1 in *Rb1^+/+^* and *Rb1^−/−^* hepatocytes from the three independent biological replicates. Each experiment used hepatocytes from different mice, and *Rb1^+/+^* and *Rb1^−/−^* hepatocytes for each replicate experiment were derived from the same mouse. The shaded area indicates the 95% confidence interval. Data were binned based on nuclear area at birth, and the mean and standard deviation of each bin is plotted. D. The R values of the linear correlations in (C) are plotted. Dashed lines link experiments performed using hepatocytes derived from the same mouse.

## Notes

### Competing Interest Statement

The authors have declared no competing interest.

